# ICP-MS analysis of metal and metalloid concentrations of common microbiological growth media reveals presence of heavy metals

**DOI:** 10.1101/2020.09.25.313221

**Authors:** Wenfa Ng

## Abstract

Trace elements such as cobalt and manganese play important roles as cofactors of enzymes, and thus, they collectively impact on biochemistry and cellular metabolism. Hence, it is of importance to gain an understanding of the trace metal and metalloid concentrations of growth medium in order to fully account for the growth performance of the bacterium. But, this aspect of microbial cell cultivation is usually neglected. Advent of instrumented techniques for metals and metalloid analysis such as inductively coupled plasma mass spectrometry (ICP-MS) has significantly facilitated trace element analysis at the parts per billion (ppb) to parts per million (ppm) level in aqueous matrixes free of organic interference. In this work, ICP-MS was utilized as the principal tool for profiling the range of metals and metalloids in different microbiological growth medium ranging from minimal salts medium to complex chemically undefined medium. Growth media examined include: LB Lennox, LB Lennox + 2 g/L glucose, LB Lennox (buffered, 89 mM phosphate), LB Lennox (buffered, 89 mM phosphate) + 6 g/L glucose, formulated medium + 6 g/L glucose, Tryptic Soy Broth, M9 medium, and M9 + 1 g/L yeast extract medium. Results reveal detection of elevated concentrations of chromium, copper and cadmium in different versions of LB Lennox and Tryptic Soy Broth. Concentrations of the above heavy metals in the tens of ppm range meant that the growth media could only support the growth of environmental bacteria with some resistance to heavy metal toxicity. On the other hand, no heavy metals were detected in M9 minimal salts medium or a modified version with supplementation of 1 g/L yeast extract. This indicated that minimal salts media may have less of a problem with heavy metal contamination compared to chemically undefined microbiological growth media. Collectively, the results highlight the essentiality of conducting a comprehensive profiling experiment for detecting different metals and metalloids at trace levels in microbiological growth medium. Such data would in addition to offering a deeper understanding of some peculiar growth behaviour from some microorganisms, may also help identify contamination issue during manufacture that preclude use of the media in cultivating many laboratory domesticated microorganisms.

**Subject areas:** biochemistry, biotechnology, cell biology, microbiology, biochemical engineering,

## 1. Introduction

Microbial growth requires carbon, nitrogen, sulphur, hydrogen, oxygen, and phosphorus. But, what is less often noted is the importance of trace elements in microbial growth and function.^1,2^ For example, zinc is an important cofactor for some enzymes.^3,4^ Naturally, the trace element requirement of different microbial species would likely be different, and this may be defined through a survey of the enzymes encoded in the genome of the organism. Besides nutritional roles of trace elements, another important aspect in understanding the elemental composition of microbiological growth medium is for quality control purposes as lapses in manufacturing may introduce unneeded contaminants into the growth medium.

The issue is particularly serious for rich medium. Typically, such media are chemically complex and undefined; thereby, leaving the scientist unable to correlate medium composition with growth behaviour of a microbe should problems of poor growth occur in the experiment. More importantly, components such as yeast extract and tryptone commonly incorporated into rich media carry an undefined elemental composition that serves as variability factors in microbial cell culture, particularly in applications such as cell differentiation studies or where the growth state of the cells are important. Indeed, yeast extract and tryptone both suffer from batch to batch variability, and this element introduces unpredictability to microbial cell culture through differences in amino acids, growth factors, vitamins, and less often noted, trace metal and metalloids compositions. Given the important role played by trace elements in serving as cofactors of enzymes and supporting other critical cellular functions,^5^ types and concentrations of trace elements in microbiological growth medium should be a facet that deserve comprehensive quantification.

Approaches for quantifying elemental compositions of microbiological medium typically revolves around instrumented approaches or those based on chemical reactions followed by calorimetric assays.^6,7^ For trace element analysis, instrumented approaches such as inductively coupled plasma mass spectrometry (ICP-MS) and inductively coupled plasma optical emission spectroscopy (ICP-OES) are suitable and go-to tools.^8-10^ Amongst the two techniques, ICP-MS is the more sensitive approach, and is able to detect, with precision, parts per billion (ppb) concentration of many metals and metalloids in the Periodic table. On the other hand, ICP-OES is a more robust and less maintenance intensive instrument for routine detection of elements in the parts per million (ppm) range in more difficult matrixes.

This work sought to quantify the presence and concentrations of a range of metals and metalloids in common microbiological growth media through ICP-MS analysis. Growth medium considered include, LB Lennox, LB Lennox + 2 g/L glucose, LB Lennox (buffered, 89 mM phosphate), LB Lennox (buffered, 89 mM phosphate) + 6 g/L glucose, formulated medium + 6 g/L glucose, Tryptic Soy Broth, M9 medium, and M9 + 1 g/L yeast extract medium. Different growth media were prepared in glass shake flasks, and the contents sterilized at 121 °C for 20 minutes in an autoclave. Sampling was done after the media have cooled to room temperature. Overall, the experimental protocol developed for this work tried to mimic the environmental conditions in the growth media as closely as possible to when it is used in actual settings for microbial cultivation.

## 2. Materials and methods

### 2.1 Chemicals and growth medium

Multi-element calibration standard for calibrating inductively coupled plasma mass spectrometry (ICP-MS) was purchased from Merck. LB medium and Tryptic Soy Broth were purchased from BD and Merck, respectively. The rest of the medium components were purchased from Sigma-Aldrich. Composition of LB Lennox medium is [g/L]: Tryptone, 10.0; Yeast extract, 5.0; NaCl, 5.0. Composition of LB Lennox + 2 g/L glucose medium is [g/L]: Tryptone, 10.0; Yeast extract, 5.0; NaCl, 5.0; D-Glucose, 2.0. Composition of LB Lennox (buffered, 89 mM phosphate) medium is [g/L]: Tryptone, 10.0; Yeast extract, 5.0; NaCl, 5.0; K_2_HPO_4_, 12.54; KH_2_PO_4_, 2.31. Composition of LB Lennox (buffered, 89 mM phosphate) + 6 g/L glucose medium is [g/L]: Tryptone, 10.0; Yeast extract, 5.0; NaCl, 5.0; D-Glucose, 6.0, K_2_HPO_4_, 12.54; KH_2_PO_4_, 2.31. Composition of formulated medium + 6 g/L glucose medium is [g/L]: K_2_HPO_4_, 12.54; KH_2_PO_4_, 2.31; D-Glucose, 4.0; NH_4_Cl, 1.0; Yeast extract, 12.0; NaCl, 5.0; MgSO_4_, 0.24; Composition of Tryptic Soy Broth is [g/L]: Peptone from casein 17.0; Peptone from soymeal 3.0; D-Glucose 2.5; NaCl 5.0; K_2_HPO_4_, 2.5. Composition of M9 medium is [g/L]: D-Glucose, 4.0; NH_4_Cl, 1.0; NaH_2_PO_4_, 3.0; Na_2_HPO_4_, 6.78; NaCl, 0.5; MgSO_4_, 0.24. Composition of M9 minimal medium supplemented with 1 g/L yeast extract was [g/L]: D-Glucose, 4.0; NH_4_Cl, 1.0; NaH_2_PO_4_, 3.0; Na_2_HPO_4_, 6.78; NaCl, 0.5; MgSO_4_, 0.24; yeast extract, 1.0.

### 2.2 Growth medium preparation and sterilization

Different growth media were constituted as is according to manufacturers’ instructions or through weighing and dissolving different medium components. The solvent used was deionized water prepared from the combined filtration and ion exchange system with cationic and anionic ion exchange resins. 100 mL media were prepared in 250 mL glass shake flask stopped with a cotton plug. After sterilization at 121 °C for 20 minutes in an autoclave, the prepared media were allowed to cool to room temperature prior to withdrawal of samples using aseptic technique in a Class II Biosafety cabinet on the day of the ICP-MS analysis.

### 2.3 Sample preparation and inductively coupled plasma mass spectrometry (ICP-MS) analysis

Using aseptic technique, 5 mL samples were obtained from each type of sterilized growth medium contained in individual shake flasks. Three samples representing three technical replicates were obtained from each type of growth medium in a single flask. Samples were diluted 100 times prior to filtration through a 0.45 μm nylon membrane filter prior to ICP-MS analysis. After filtration, samples were placed in 20 mL polyethylene sample bottle.

Agilent 7500a ICP-MS was calibrated by a calibration series of 0, 50 ppb, 100 ppb, 500 ppb, 1000 ppb, 5000 ppb, 10000 ppb calibration standards obtained from diluting a Merck multi-element calibration standard with 18.2 MΩ ultrapure water. Instrument was operated under hot plasma mode (1250 W). Fresh calibration standards were used for calibrating the instrument. Detection was attempted for each calibrated element and isotope in each sample.

## 3. Results and Discussion

**Table 1:** Metals and metalloids concentration in LB Lennox medium.

| **Elements** | **Concentration / ppm** |
| --- | --- |
| <sup>7</sup>Li | 59.07 |
| <sup>11</sup>B | 12.60 |
| <sup>24</sup>Mg | 197.87 |
| <sup>27</sup>Al | 6.02 |
| <sup>43</sup>Ca | 202.87 |
| <sup>53</sup>Cr | 75.15 |
| <sup>55</sup>Mn | 1.17 |
| <sup>56</sup>Fe | 374.00 |
| <sup>57</sup>Fe | 1331.00 |
| <sup>59</sup>Co | 0.04 |
| <sup>60</sup>Ni | 1.77 |
| <sup>63</sup>Cu | 16.62 |
| <sup>66</sup>Zn | 0.89 |
| <sup>69</sup>Zn | 0.00 |
| <sup>106</sup>Cd | 13.71 |
| <sup>108</sup>Cd | 0.00 |
| <sup>111</sup>Cd | 0.00 |
| <sup>137</sup>Ba | 0.41 |
| <sup>206</sup>Pb | 0.00 |
| <sup>207</sup>Pb | 0.00 |
| <sup>208</sup>Pb | 0.00 |
| <sup>209</sup>Bi | 1.90 |

**Table 2:**
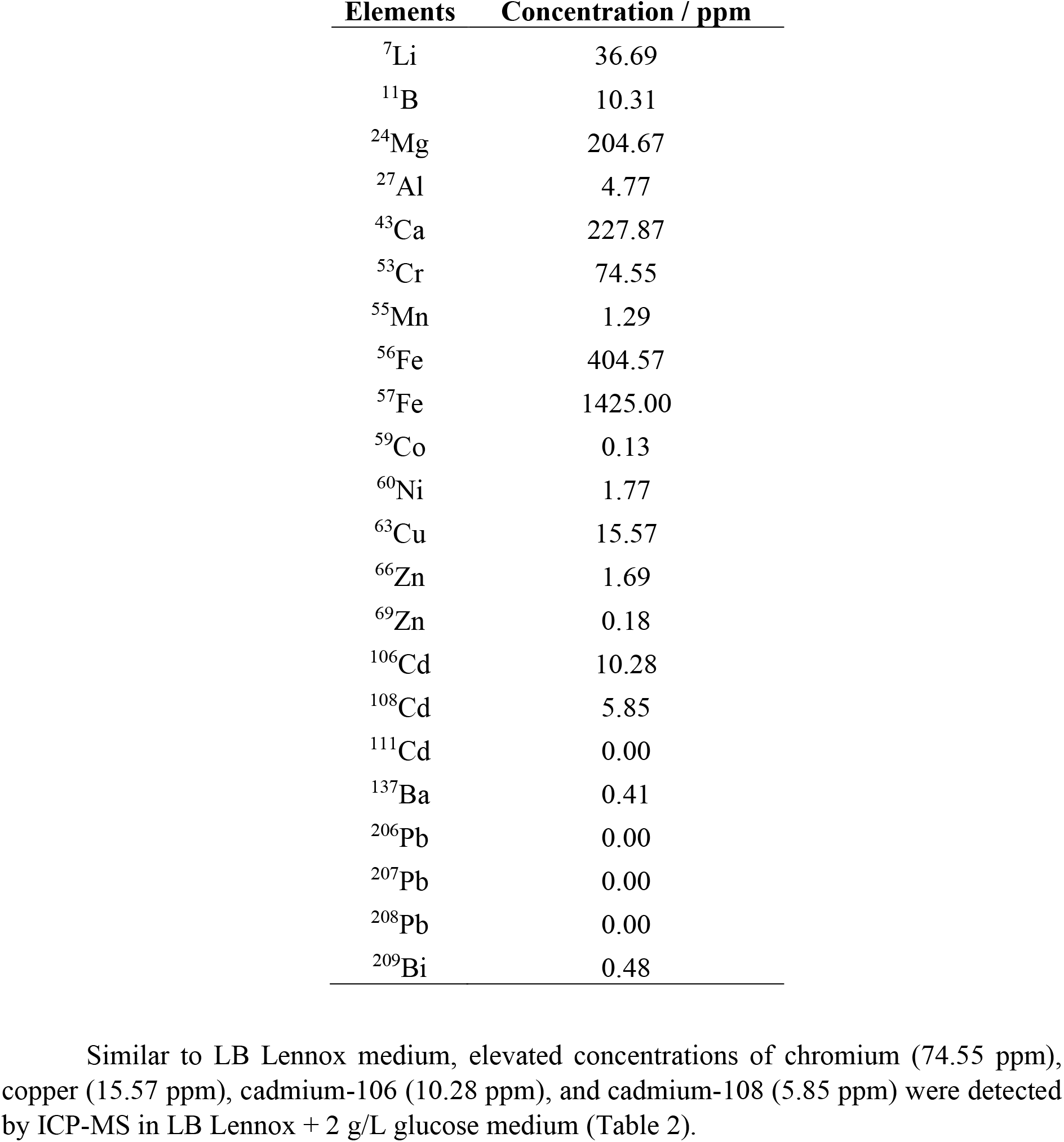
Metals and metalloids concentration in LB Lennox + 2 g/L glucose medium.

**Table 3:** Metals and metalloids concentration in LB Lennox (89 mM phosphate) medium.

| **Elements** | **Concentration / ppm** |
| --- | --- |
| <sup>7</sup>Li | 5.39 |
| <sup>11</sup>B | 14.61 |
| <sup>24</sup>Mg | 156.47 |
| <sup>27</sup>Al | 4.42 |
| <sup>43</sup>Ca | 177.87 |
| <sup>53</sup>Cr | 84.77 |
| <sup>55</sup>Mn | 1.79 |
| <sup>56</sup>Fe | 413.40 |
| <sup>57</sup>Fe | 1157.00 |
| <sup>59</sup>Co | 0.06 |
| <sup>60</sup>Ni | 2.52 |
| <sup>63</sup>Cu | 36.15 |
| <sup>66</sup>Zn | 0.53 |
| <sup>69</sup>Zn | 0.00 |
| <sup>106</sup>Cd | 0.00 |
| <sup>108</sup>Cd | 0.00 |
| <sup>111</sup>Cd | 0.00 |
| <sup>137</sup>Ba | 0.41 |
| <sup>206</sup>Pb | 0.00 |
| <sup>207</sup>Pb | 0.00 |
| <sup>208</sup>Pb | 0.00 |
| <sup>209</sup>Bi | 0.11 |

**Table 4:** Metals and metalloids concentration in LB Lennox (89 mM phosphate) + 6 g/L glucose medium.

| **Elements** | **Concentration / ppm** |
| --- | --- |
| <sup>7</sup>Li | Not detected |
| <sup>11</sup>B | 31.34 |
| <sup>24</sup>Mg | 126.17 |
| <sup>27</sup>Al | 15.38 |
| <sup>43</sup>Ca | 101.47 |
| <sup>53</sup>Cr | 22.53 |
| <sup>55</sup>Mn | 227.47 |
| <sup>56</sup>Fe | 255.90 |
| <sup>57</sup>Fe | 945.30 |
| <sup>59</sup>Co | 0.65 |
| <sup>60</sup>Ni | 3.19 |
| <sup>63</sup>Cu | 46.83 |
| <sup>66</sup>Zn | 0.00 |
| <sup>69</sup>Zn | 0.06 |
| <sup>106</sup>Cd | 0.00 |
| <sup>108</sup>Cd | 0.00 |
| <sup>111</sup>Cd | 1.40 |
| <sup>137</sup>Ba | 0.15 |
| <sup>206</sup>Pb | 0.41 |
| <sup>207</sup>Pb | 0.22 |
| <sup>208</sup>Pb | 0.00 |
| <sup>209</sup>Bi | 0.62 |

**Table 5:** Metals and metalloids concentration in formulated medium + 6 g/L glucose medium.

| **Elements** | **Concentration / ppm** |
| --- | --- |
| <sup>7</sup>Li | 0.00 |
| <sup>11</sup>B | 12.62 |
| <sup>24</sup>Mg | 189.93 |
| <sup>27</sup>Al | 3.49 |
| <sup>43</sup>Ca | 170.63 |
| <sup>53</sup>Cr | 77.68 |
| <sup>55</sup>Mn | 1.82 |
| <sup>56</sup>Fe | 388.47 |
| <sup>57</sup>Fe | 1017.63 |
| <sup>59</sup>Co | 0.00 |
| <sup>60</sup>Ni | 2.70 |
| <sup>63</sup>Cu | 42.21 |
| <sup>66</sup>Zn | 0.89 |
| <sup>69</sup>Zn | 0.00 |
| <sup>106</sup>Cd | 17.14 |
| <sup>108</sup>Cd | 2.92 |
| <sup>111</sup>Cd | 0.00 |
| <sup>137</sup>Ba | 0.41 |
| <sup>206</sup>Pb | 0.00 |
| <sup>207</sup>Pb | 0.00 |
| <sup>208</sup>Pb | 0.00 |
| <sup>209</sup>Bi | 0.58 |

**Table 6:** Metals and metalloids concentration in Tryptic soy broth.

| Elements | Concentration / ppm |
| --- | --- |
| <sup>7</sup>Li | 117.39 |
| <sup>11</sup>B | 21.52 |
| <sup>24</sup>Mg | 162.83 |
| <sup>27</sup>Al | 9.58 |
| <sup>43</sup>Ca | 195.57 |
| <sup>53</sup>Cr | 64.21 |
| <sup>55</sup>Mn | 1.48 |
| <sup>56</sup>Fe | 325.03 |
| <sup>57</sup>Fe | 1011.47 |
| <sup>59</sup>Co | 0.52 |
| <sup>60</sup>Ni | 1.21 |
| <sup>63</sup>Cu | 30.55 |
| <sup>66</sup>Zn | 3.20 |
| <sup>69</sup>Zn | 0.71 |
| <sup>106</sup>Cd | 6.85 |
| <sup>108</sup>Cd | 8.77 |
| <sup>111</sup>Cd | 0.00 |
| <sup>137</sup>Ba | 1.23 |
| <sup>206</sup>Pb | 2.95 |
| <sup>207</sup>Pb | 1.43 |
| <sup>208</sup>Pb | 1.92 |
| <sup>209</sup>Bi | 4.96 |

**Table 7:**
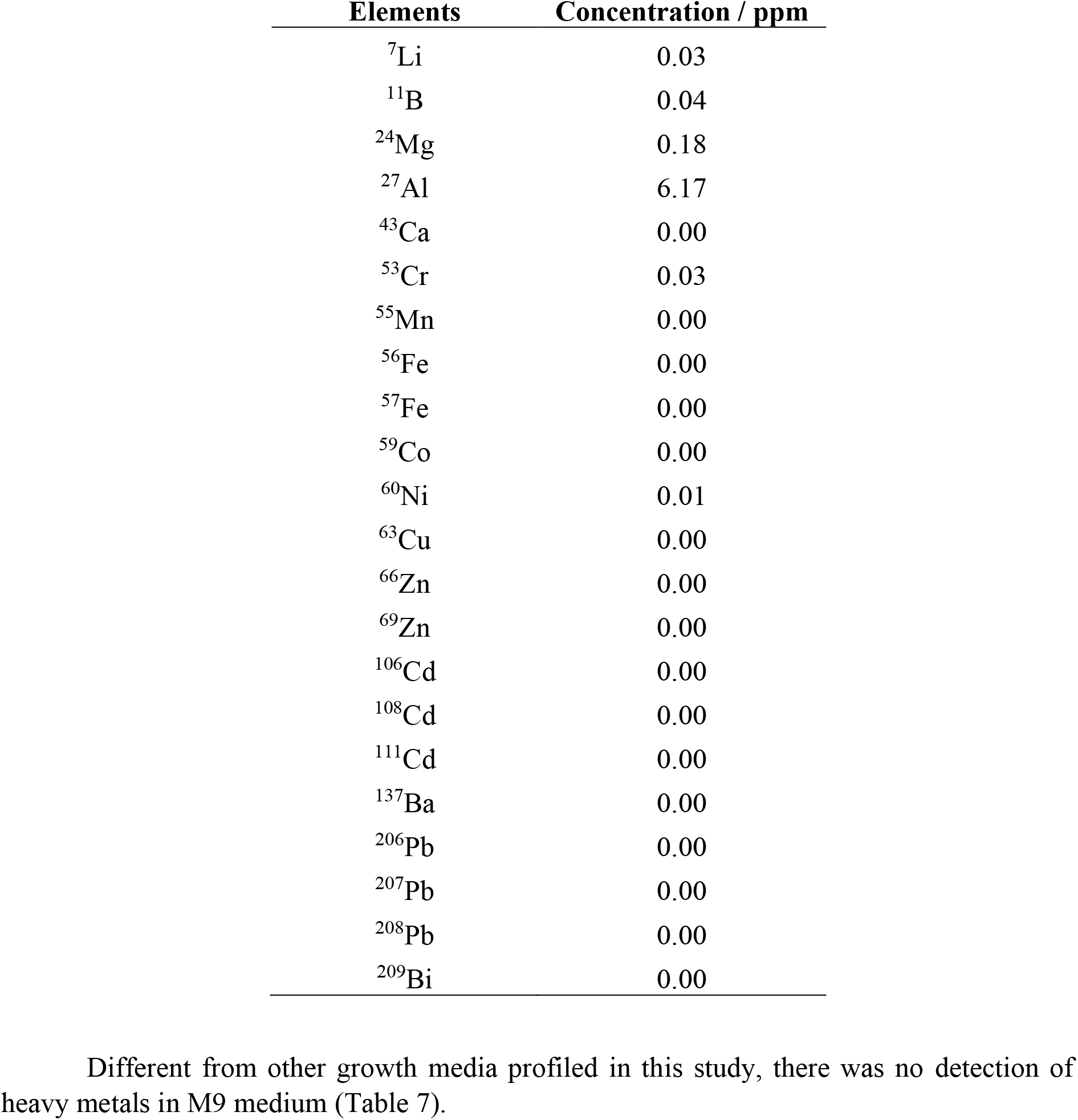
Metals and metalloids concentration in M9 medium.

**Table 8:**
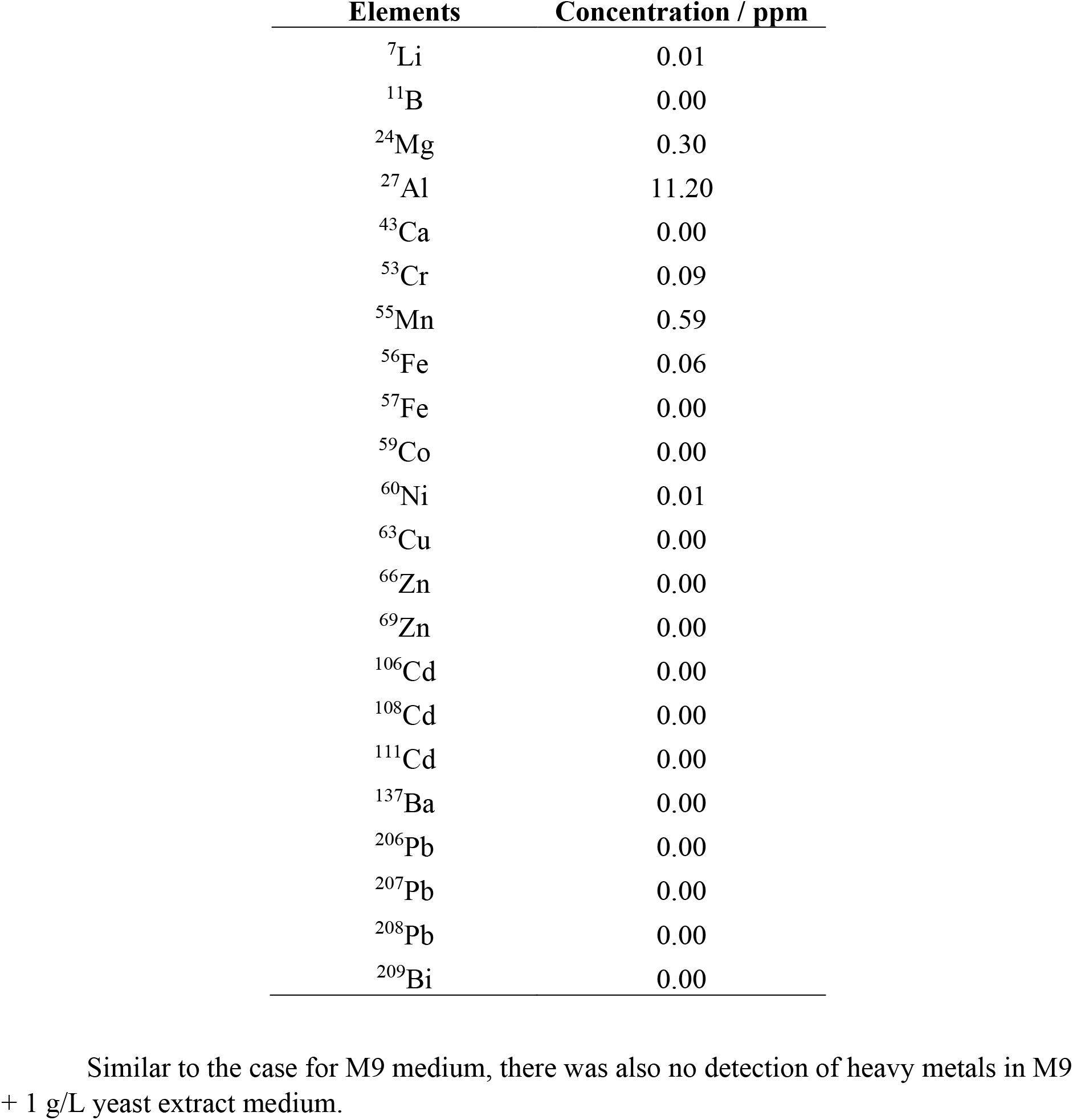
Metals and metalloids concentration in M9 + 1 g/L yeast extract medium.

## 4. Conclusions

Trace elements such as metals and metalloids play important roles in biochemistry and cellular metabolism. Thus, their provision in growth medium is critical for ensuring good growth performance from microorganisms. This study sought to investigate the trace metal element concentrations in a variety of common microbiological media ranging from minimal salts to complex chemically undefined medium. Results reveal unexpected elevated concentrations of heavy metals in many complex undefined media such as LB Lennox and Tryptic Soy Broth. Heavy metals of concern are principally chromium, copper, cadmium, and lead. Concentrations of heavy metals in the tens of parts per million meant that the media could only be used in cultivation of bacteria bearing some resistance to heavy metal toxicity effect.

This then limits the utility of the growth media to cultivation of environmental microbes as compared to laboratory domesticated species. On the contrary, no heavy metals were detected in minimal salts medium such as M9 medium. Overall, the results of this study highlight the importance of gaining understanding of the metal and metalloid concentrations in different growth media both from the perspective of quality control purposes as well as understanding growth performance of particular microbe in the medium.

